# *Pediococcus pentosaceus* decreases α-synuclein accumulation and alleviates parkinsonism in *Caenorhabditis elegans* and *Oryzias latipes*

**DOI:** 10.1101/2025.01.17.633680

**Authors:** Tomomi Komura, Masayuki Yoshida, Rio Kurihara, Masato Kinoshita, Masaru Yoshida, Yoshikazu Nishikawa

**Affiliations:** Graduate School of Human Science and Environment, University of Hyogo, 1-1-12 Shinzaike-Honcho, Himeji, Hyogo 670-0092, Japan; Research Institute for Food and Nutritional Sciences, University of Hyogo, 1-1-12 Shinzaike-Honcho, Himeji, Hyogo 670-0092, Japan; Graduate School of Integrated Sciences for Life, Hiroshima University, 1-4-4 Kagamiyama, Higashihiroshima 739-8528, Japan; Division of Applied Bioscience, Graduate School of Agriculture, Kyoto University, Kitashirakawa-Oiwake-cho, Sakyo-ku, Kyoto 606-8502, Japan; Faculty of Food and Nutrition Science, Tezukayama Gakuin University, 4-2-2 Harumidai, Minami-ku Sakai, Osaka 590-0113, Japan

**Keywords:** *C. elegans*, *O. latipes*, Parkinson’s disease, α-synuclein, lactic acid bacteria, *P. pentosaceus*

## Abstract

In Parkinson’s disease (PD), α-synuclein (αSyn) accumulation drives neuropathological progression, establishing it as a potential therapeutic target. Lactic acid bacteria (LAB) provide various health benefits to the host and are expected to offer protective effects on neurological functions through the brain–gut connection, as indicated by animal studies. However, the protective effects of LAB against αSyn are not well understood. We investigated whether LAB feeding could reduce αSyn accumulation and improve mobility in transgenic *Caenorhabditis elegans*, an invertebrate model organism that expresses human αSyn in muscle. Among the nine screened strains, *Pediococcus pentosaceus* suppressed αSyn accumulation and the decrease in bending counts, a locomotion index in nematodes. Additionally, feeding *P. pentosaceus* to *Oryzias latipes* (medaka), a vertebrate model organism, alleviated PD-like behavioral defects induced by 1-methyl-4-phenyl-1,2,3,6-tetrahydropyridine. *P. pentosaseus* acts as a probiotic against PD, and PD model worms and medaka can be used to screen effective LAB for prevention.

## 1. Introduction

Parkinson’s disease (PD) is the second most common neurodegenerative disease, after Alzheimer’s disease, and the most common disease that causes movement disorders (Dorsey, Sherer, Okun, & Bloem, 2018). Braak et al. indicated that α-synuclein (αSyn) deposits are associated with disease progression (Braak et al., 2003). As the magnitude of αSyn accumulation affects symptom progression and expansion of the pathological lesion in the brain, αSyn is now considered a potential therapeutic target (Menon et al., 2022); thus, its accumulation must be prevented at an early stage (Mochizuki, Choong, & Masliah, 2018).

Phenolic metabolites digested and elevated in the gut microbiota of mice could inhibit αSyn aggregation (Ho et al., 2019). These metabolites were also effective in inhibiting the progression of motility dysfunction owing to αSyn aggregation in a fruit fly model (Ho et al., 2019). Therefore, the gut microbiota seems to be involved in the regulation of αSyn aggregation and PD-like symptoms (Zhang, Tang, & Guo, 2023). Indeed, lactic acid bacteria (LAB) prevented αSyn accumulation and improved motor behavior and neuroinflammation in mouse models (Perez Visnuk, Savoy de Giori, LeBlanc, & de Moreno de LeBlanc, 2020; Wang et al., 2022). However, it takes several months or longer to examine the effects of LAB on αSyn aggregation in mice.

*Caenorhabditis elegan* (nematode) is a model organism used in genetic experiments since 1974 (Brenner, 1974). Molecular biology techniques have been established in this species, making it a useful host model for aging and host-microbe studies (Kim & Flavell, 2020). Alzheimer’s disease is the most common age-related neurological disease caused by the accumulation of the pathogenic protein amyloid-β in the brain, which induces nerve cell degeneration and dementia (Haass & Selkoe, 2007). Previously, we found that *Pediococcus parvulus* and *Lactococcus laudensis* reduced amyloid-β in *C. elegans* (Komura, Aoki, Kotoura, & Nishikawa, 2022). It seems possible to find an effective LAB for the prevention of PD using nematodes. PD models have also been produced using *Danio rerio* (zebrafish) or *Oryzias latipes* (medaka) as vertebrates (Doyle & Croll, 2022;,Uemura et al., 2015). However, to date, the preventive effects of LAB on PD have not been reported.

This study screened LAB of the *Pediococcus* and *Lactococcus* species for their ability to suppress αSyn aggregation in *C. elegans*. Afterward, the efficacy of LAB in alleviating PD-like symptoms was evaluated using a larval model of medaka, as the medaka genome size is 800 Mbp, half that of zebrafish (Naruse, Chisada, Sasado, & Takehana, 2016), and would be easier for genetic analysis.

## 2. Material and methods

### 2.1 Maintenance of nematodes and medaka

Wild type (Bristol N2) and transgenic nematode strains were provided by the Caenorhabditis Genetics Center of the University of Minnesota. The transgenic worm used in this study was NL5901, *unc-54p::alphasynuclein::YFP + unc-119*. The NL5901 strain exhibits αSyn expression in the muscles. Nematodes were maintained at 25 °C and propagated on nematode growth medium according to standard techniques. Cultured bacteria (100 mg wet weight) were suspended in 0.5 mL M9 buffer (3 g KH_2_PO_4_, 6 g Na_2_HPO_4_, 5 g NaCl, 1 mL 1 M MgSO_4_, H_2_O to 1 L), and 50 μL of the resulting bacterial suspension was then spread on peptone-free nematode growth medium (pfNGM) plates (5.5 cm diameter) to feed the worms (Hoshino et al., 2008).

An inbred medaka line, Cab, was maintained at 27 °C under a 14-h light/10-h dark cycle in a recirculating aquaculture system equipped with a carbon filtration system, ultraviolet light sterilizers, and a biofiltration system (Iwaki Aquatic Co Ltd., Tokyo, Japan). Embryos were collected via natural spawning and raised in breeding water containing 0.0001% methylene blue until hatching.

### 2.2 Ethical statement

All experimental protocols were approved by the Institutional Animal Care and Use Committee of the University of Hyogo (Approval No. 017, dated April 20, 2023). Tricaine was used to anesthetize animals when necessary. The medaka experiment was conducted in accordance with ARRIVE guidelines. Since this study focused on post-hatching larvae, sex differences were not assessed.

### 2.3 Bacterial strains

*E. coli* OP50 (OP50) is an internationally established food for nematodes. Tryptone soya agar (Nissui Pharmaceutical, Tokyo, Japan) was used to culture OP50 cells. LAB were cultured aerobically using De Man–Rogosa–Sharpe broth (Becton Dickinson, New Jersey, USA) at 30 °C for 48 h (Table 1).

**Table 1.** Lactic acid bacteria used in this study.

| Bacterial strains | Isolation source | Source |
| --- | --- | --- |
| *Lactococcus raffinolactis* NBRC100932 | Raw milk | NBRC |
| *Lactococcus plantarum* NBRC100936 | Frozen peas | NBRC |
| *Lactococcus garvieae* strain MCRI-184 | Cured meat | (Komura et al., 2022) |
| *Lactococcus lactis subsp. lactis* MCRI- 595 | Gingers | (Komura et al., 2022) |
| *Lactococcus laudensis* MCRI-603 | Kales | (Komura et al., 2022) |
| *Pediococcus damnosus* NBRC107867 | Lager beer yeast | NBRC |
| *Pediococcus acidilactici* NBRC109619 | Barley | NBRC |
| *Pediococcus parvulus* MCRI-590 | Juice from Japanese turnip<br>pickles | (Komura et al., 2022) |
| *Pediococcus pentosaceus* MCRI-640 | Purple sweet potato | This study |

NBRC, NITE Biological Resource Center; MCRI, Marudai Central Research Institute.

LAB isolated from food were provided by the Marudai Central Research Institute (MCRI) of Marudai Foods Co. (Osaka, Japan) or NITE Biological Resource Center. MCRI strains were isolated and identified as reported previously (Komura, Aoki, & Nishikawa, 2024).

### 2.4 Measurement of αSyn expression in nematodes

The nematodes were maintained according to the international standard method and in pfNGM plates, as previously described (Komura, Ikeda, Yasui, Saeki, & Nishikawa, 2013). All assays were performed by adding young adult worms (3 days old) to each pfNGM plate (5.5 cm in diameter) covered with LAB or OP50 at 25 °C.

After treatment with OP50 or LABs for 24 h from young adults (strain NL5901), each worm was washed three times with M9 buffer to eliminate adhering bacteria and immobilized at a final concentration of 50 mM sodium azide. After mounting on 5% agar pad slides made with M9 buffer containing 10 mM sodium azide and sealing the coverslips, images were recorded using an upright fluorescence microscope (BX61; Olympus Co., Tokyo, Japan) with a 40× objective lens (U Plan FL N 40×/0.75) and a USB camera WRAYCAM-VEX830 (WRAYMER Inc., Osaka, Japan). The measured YFP values were normalized to the density values determined for the head size from the mouth to the pharynx. Next, YFP values were measured using ImageQuant TL version 8 (Cytiva, Tokyo, Japan), and Adobe Photoshop Elements were normalized to density values per square millimeter of the projection area of the worm using ImageJ v1.54 software developed by the National Institutes of Health. The assay was performed using more than 10 worms for analysis and two biological replicates.

### 2.5 Determination of nematode motility based on body bends

Young adult worms (3 days old) of wild type and the NL5901 strain were treated with LAB or OP50 for 7 days at 25 °C. The nematodes (10 days old) were collected and transferred in a drop of M9 buffer onto pfNGM, allowed to sit for 10 s, and the number of body bends of individual nematodes was counted microscopically for 1 min using a STZ-161-TLED stereomicroscope (Shimadzu Co. Ltd., Kyoto, Japan) and a USB camera. A body bend was defined as a change in the direction of midbody bending. The number of body bends was counted for 1 min. The assay was performed using at least 35 worms for analysis and two biological replicates.

### 2.6 Swimming activity of medaka larvae

For medaka, immediately after hatching, 10 larvae were sorted and placed into plastic plates (5.5 cm in diameter) in 10 mL of Yamamoto’s Ringer solution [0.75% NaCl, 0.02% KCl, 0.02% CaCl_2_, 0.002% NaHCO_3_ (adjusted to pH 7.3 with 5% NaHCO_3_)] (Murata, Kinoshita, Naruse, Tanaka, & Kamei, 2019). The solutions were replaced daily with fresh ones. A solution of 1-methyl-4-phenyl-1,2,3,6-tetrahydropyridine (MPTP) hydrochloride (M2690, Tokyo Chemical Industry Co., Ltd., Tokyo, Japan) was prepared using Yamamoto’s ringer. A suitable MPTP concentration was determined based on the results of Matsui et al. (Matsui et al., 2009). For medaka feeding, LAB were recovered from the De Man–Rogosa–Sharpe broth and washed with phosphate-buffered saline, after which the bacterial sediment was dried on a clean bench. The dried LAB were stored at -80 °C until fed to medaka and ground finely with a needle before feeding. Medaka larvae were fed twice daily immediately after hatching in sufficient amounts.

The locomotor activities of the control and LAB- and/or MPTP-treated larvae were measured individually. To confirm that they were eating the food (Otohime B1, Marubeni Nisshin Feed Co., Ltd., Tokyo, Japan), they were fed at different times twice a day for 15 days. Exposure to MPTP was performed for 2 d, beginning 13 d after hatching. The videos were analyzed as mentioned below.

First, 15-day-old larvae were subjected to the tests at room temperature (22 °C). There were no statistically significant differences in body size between the groups. Then, a larva was gently introduced into a plastic cuvette containing Yamamoto’s Ringer solution, and a 5-min video recording from the side was then started using a digital camera (TG-6, Olympus, Tokyo, Japan) at 60 frames/s after settling for 2 min. The cuvette was illuminated from the back by using a diffuser composed of a white acrylic plate to enhance the contrast of the video images.

The video was downsampled to 20 frames/s, and the positions of the larvae in all video frames were tracked using the video tracking software DeepLabCut (Nath et al., 2019). The total traveling distance and median vertical position of the larva in the 5-min recording period were calculated from the obtained coordinates as measures of motor performance.

### 2.7 Statistical analysis

The αSyn expression, body bending counts in worms, and swimming activity for medaka larvae were compared using the nonparametric Steel–Dwass method. Where significant differences were observed, the data were classified as * *p* < 0.05 and *\*\*p* < 0.01. All statistical analyses were performed using Microsoft Excel supplemented with the add-in software + Statcel 3 (OMS, Tokyo, Japan).

## 3. Results and discussion

To observe the expression and aggregation of αSyn, we used the *C. elegans* strain NL5901, constructed with human αSyn fused to yellow fluorescent protein (YFP) under the control of the *unc-54* promoter, whose gene expression occurred in the body wall muscle cells (van Ham et al., 2008). The advantages of this strain are its high expression efficiency and the PD-like progressive defects of *C. elegans* motility (Cooper et al., 2015); thus, it demonstrates the *in vivo* toxicity of aggregation. First, we analyzed *unc-54p*::YFP expression in the entire body of *C. elegans*. Worms fed *P. pentosaceus, P. acidilactici*, or *L. plantarum* showed significantly lower fluorescence intensity than those fed *E. coli* OP50 or the other six LAB strains (Fig. 1A-D). YFP assembly showed αSyn aggregation, and the particles were fewer in worms fed with the three strains (Fig. 1A, B); thus, the expression and/or aggregation of αSyn seemed to be suppressed.

**Fig. 1.**
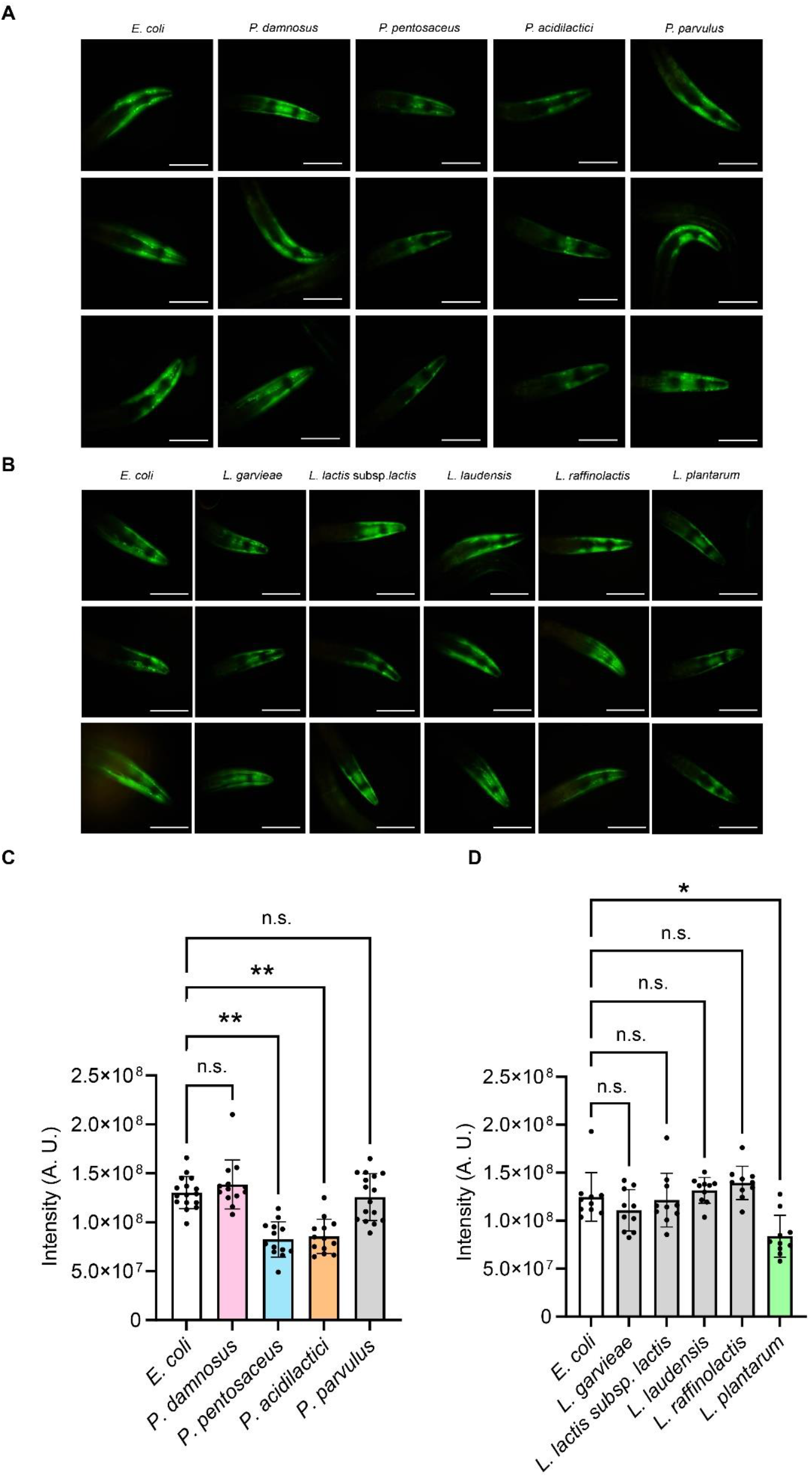
Yellow fluorescent protein (YFP) signals of the α-synuclein expression in the NL5901 strain of *C. elegans* treated with OP50 or *Pediococcus* (A) or *Lactococcus* (B). Scale bar, 0.2 mm. Graphical representation showing the quantification of YFP-tagged α-synuclein in NL5901. The data represent the mean fluorescent intensity with SD. Each point represents a single worm assessed. Asterisks * and ** indicate statistically significant differences at *p* < 0.05 and < 0.01, respectively (C and D).

The body bending counts of the transgenic strain NL5901 were significantly lower than those of the wild-type N2 at 10 days old; however, several NL5901 worms fed *P. pentosaceus* and *P. acidilactici* showed significant recovery of bending counts (Fig. 2). PD causes movement disorders, such as shakiness of the limbs and rigidity(Yang et al., 2023). Although the mechanism is still unknown, it was reported that αSyn aggregation causes muscle atrophy owing to neuromuscular junction degeneration (Yang et al., 2023). Hence, elderly patients with PD are more susceptible to frailty and sarcopenia when severely affected (Vetrano et al., 2018). In the advanced stages of PD, it is important to screen for frailty because of the risk of falls, and to assemble a treatment regimen that maintains independence in daily life activities (Drey et al., 2017). If certain LAB are capable of inhibiting αSyn, their intake could prevent frailty and PD throughout daily life. As *P. pentosaceus* and *P. acidilactici* suppressed αSyn, these strains are expected to prevent frailty owing to its accumulation. Hence, we examined whether the bacteria could be effective in vertebrate PD models.

**Fig. 2.**
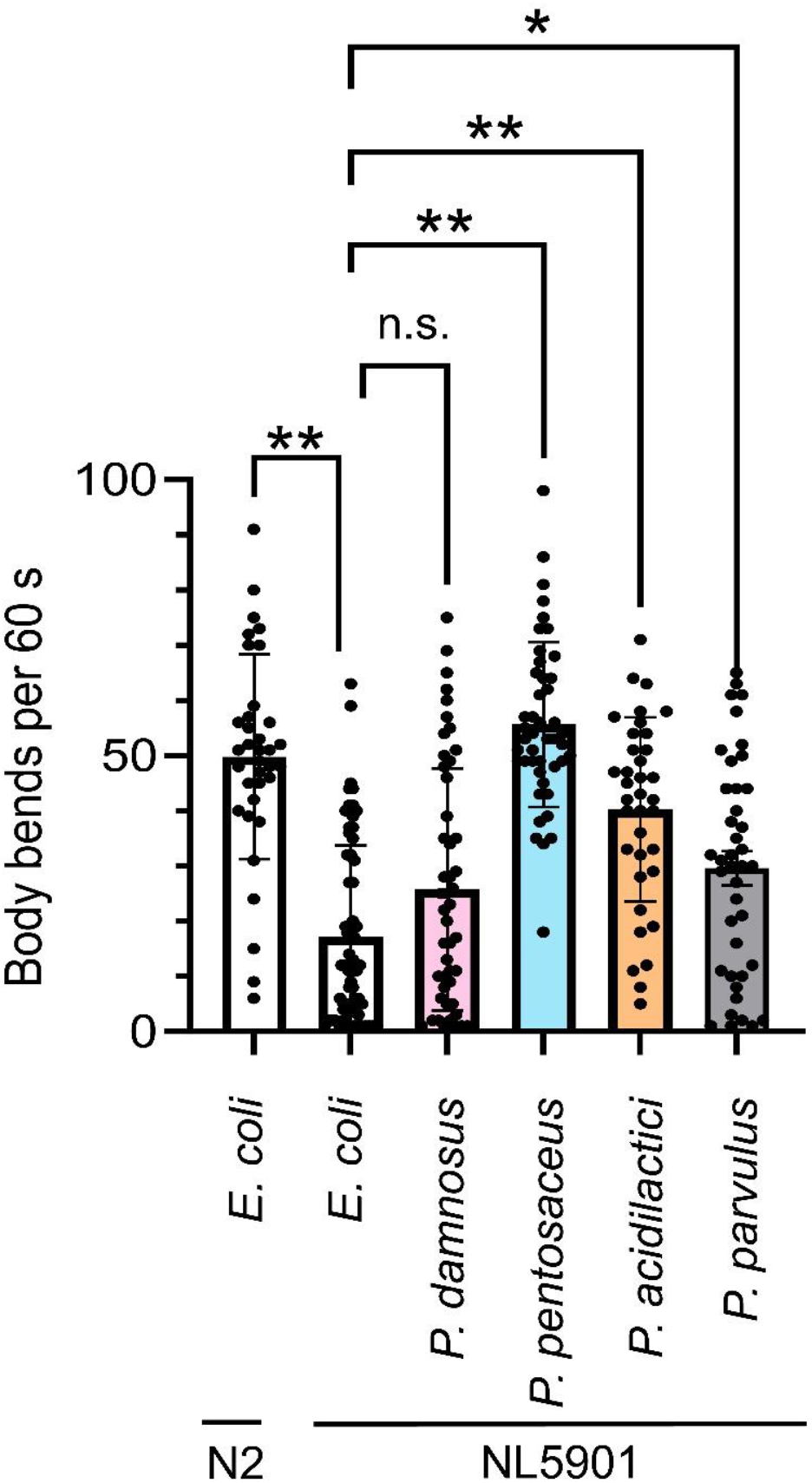
Count numbers of body bends per 1 min were assessed in N2 and NL5901 nematode strains in *E. coli* OP50 in the control or *Pediococcus* genus. The data represent the mean counts of body bends with SD. Each point represents a single worm assessed. Asterisks * and ** indicate statistically significant differences at *p* < 0.05 and < 0.01, respectively.

MPTP causes the loss of dopaminergic neurons and results in Parkinson-like symptoms of movement and behavioral abnormalities in a wide variety of vertebrates (Jackson-Lewis & Przedborski, 2007; Jakowec & Petzinger, 2004; Matsui et al., 2009). Medaka larvae fed *P. pentosaceus*, which effectively inhibited αSyn expression and recovered bending counts in *C. elegans*, showed resistance against MPTP treatment, whereas the control larvae and those fed *P. damnosus* swam less and rested at the bottom (Fig. 3A). Swimming distance and vertical movement values over a 5-min period were significantly reduced by MPTP, except in worms fed *P. pentosaceus* (Fig. 3B, C).

**Fig. 3.**
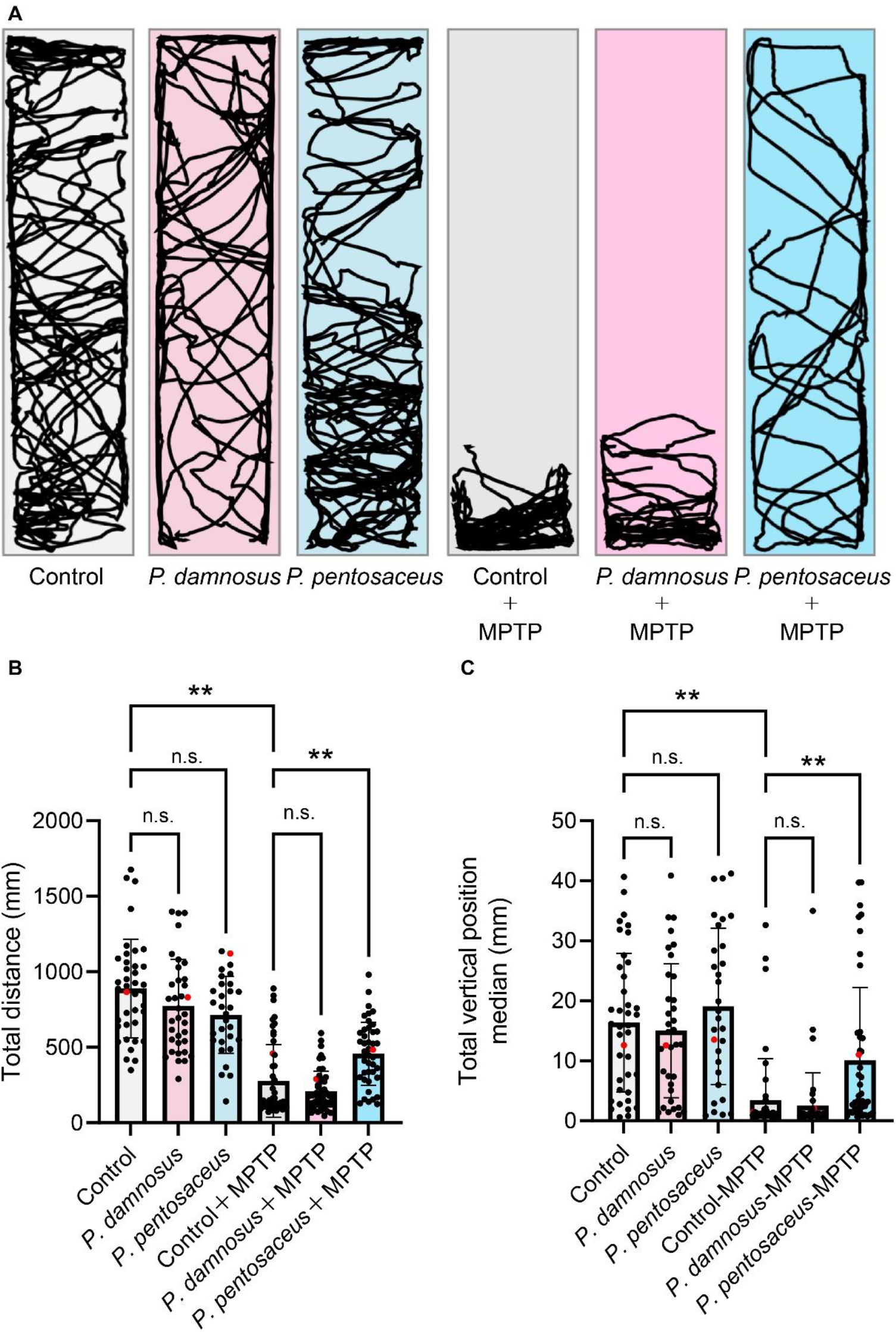
Swimming activity of medaka larvae in 1-methyl-4-phenyl-1,2,3,6-tetrahydropyridine (MPTP) treatment. The larvae were fed *Pediococcus pentosaseus* or *Pediococcus damnosus* before MPTP treatment. The tracking images indicate a side view of the cuvette (A). The column graph represents the mean distance (B) and vertical distance (C) traveled by the larvae in 5 min with SE. Each point represents a single larva assessed. The red points on each bar indicate the data shown in Figure 3A. An asterisk (**) indicates statistically significant differences between the control without LAB and each *Pediococcus* strain (*p* ≤ 0.01).

Recently, Pan et al. reported that feeding *P. pentosaceus*, a therapeutic agent, ameliorated MPTP-induced motor deficits in mice (Pan et al., 2022). In contrast, our study suggested a preventive effect of *P. pentosaceus*, which was administered for 2 weeks before MPTP treatment. *P. pentosaceus* is effective against Parkinson-like symptoms in worms, medaka, and mice. Oxidative stress is also involved in the pathogenesis of PD (Dias, Junn, & Mouradian, 2013). *P. pentosaceus* regulates the levels of Nrf2 (a transcription factor important for cellular responses to oxidative stress) pathway-related proteins in the brain of MPTP-induced mice (Pan et al., 2022). Although the mechanisms whereby *P. pentosaceus* prevents PD have not been clarified in the present study, *C. elegans* and *O. latipes* have been known to harbor genes homologous to Nrf2 (Loboda, Damulewicz, Pyza, Jozkowicz, & Dulak, 2016;,Nakayama et al., 2020). Analyses using knockout mutants, gene knockdown, or genome-editing techniques could validate whether Nrf2 plays a role in PD prevention in worm and medaka models. Most strains of *P. pentosaceus* produce gamma amino butyric acid (GABA) (Jiang, Cai, Lv, & Li, 2021). Reports have shown that GABA can alleviate PD symptoms (Song et al., 2021) and that it possibly modulates brain function through the intestinal tract. In the future, the active components of *P. pentosaceus* could be identified by analyzing metabolites of *P. pentosaceus*, including GABA.

Most experiments using mammals, such as mice, have examined only one type of LAB strain; however, multiple strains can be easily compared using nematodes and medaka. In this study, we compared nine LAB strains from a worm model and two strains from a medaka model. Our approach could be useful for the comparative analysis of various strains before extrapolation to mammalian models.

## 4. Conclusions

This is the first report to show that lactic acid bacteria could be preventative in *C. elegan*s and medaka (vertebrate) PD models. Although the mechanisms whereby *P. pentosaceus* prevents PD have not been clarified, this approach could be useful to compare various probiotic strains before extrapolation to mammalian models.

## Declarations of interest

None.

## Author contributions

**Tomomi Komura:** conceptualization (lead), investigation (lead), writing – original draft (lead), **Masayuki Yoshida, Masato Kinoshita, Rio Kurihara, Masaru Yoshida:** investigation (supporting), **Yoshikazu Nishikawa:** writing – original draft (supporting). All authors read and approved the final version of this article.

## Data availability

The data sets generated in this study are available from the corresponding author upon request.

## Acknowledgments

We are deeply grateful to Marudai Co., Ltd. for providing the LAB isolated from food products. The nematodes used in this study were provided by the Caenorhabditis Genetics Center, funded by the NIH Office of Research Infrastructure Programs (P40 OD010440). We would like to express our sincere thanks to Ms. Yasuna Seki at the School of Human Science and Environment, University of Hyogo, for her technical support with the swimming activity of the medaka larvae. We thank Editage (www.editage.com) for English language editing.

## Funding

This work was supported by a Grant-in-Aid for Scientific Research (C) from the Japan Society for the Promotion of Science (T.K.) [22K11781] and the TOYO SUISAN Foundation (FY2023).

## Notes

### Competing Interest Statement

The authors have declared no competing interest.

